# “A chromosome-level genome assembly for the colonial marine hydrozoan *Podocoryna americana*”

**DOI:** 10.64898/2026.03.04.709628

**Authors:** E. Sally Chang, Michael T. Connelly, Matthew Travert, Sofia N. Barreira, Alberto M. Rivera, Amanda M. Katzer, Reynold Yu, Paulyn Cartwright, Andreas D. Baxevanis

## Abstract

Cnidarians are important models for the studying the evolution of animal development, regeneration, cell type differentiation, and allorecognition. The marine hydrozoan *Podocoryna americana* is related to the well-established model species *Hydractinia symbiolongicarpus*. Although both species possess a sessile polyp stage, *P. americana* differs in that it also has a free-swimming medusa (jellyfish) stage in its life cycle. We used a combination of PacBio CLR long-read and Illumina Hi-C short-read genome sequencing to produce a chromosome-level genome assembly for *P. americana*. The final assembly is 327 Mbp in total length with 17 chromosome-scale scaffolds representing 98% of the assembly. Comprehensive functional annotation with BRAKER3 generated a total of 19,085 predicted protein-coding genes in this assembly, covering 91.2% of the metazoan BUCSO gene set. Comparison of the *P. americana* genome to other chromosome-level cnidarian genome assemblies revealed a high degree of macrosynteny conservation, and ortholog identification and gene family evolution analysis identified 522 expanded and 1,026 contracted gene families in *P. americana*. This high-quality, chromosome-level genome assembly of *P. americana* will be an invaluable resource for researchers studying the evolution of development, regeneration, and allorecognition in cnidarians and other metazoans.

## Introduction

The renewed interest in the genomes and biology of non-bilaterian animals has contributed to our understanding of the vast diversity and complexity of molecular innovations observed in the early evolution of animals. Cnidaria, a non-bilaterian phylum characterized by their use of cnidocytes for prey capture and predator defense, are of particular interest given their phylogenetic position as the sister group to Bilateria. The study of animals within this phylum, which includes corals, sea anemones, jellyfish, and hydroids, have significantly advanced our knowledge of the mechanisms underlying numerous biological processes, most notably those involved in regeneration (Gahan et al. 2016; Layden et al. 2016; Juliano et al. 2014), senescence (Bellantuono et al. 2015), neurobiology (Bosch et al. 2017; Kelava et al. 2015), and allorecognition (Rosengarten and Nicotra 2011; Nicotra 2019). Despite their early diverging phylogenetic position, cnidarian genomes are remarkably similar to the human genome in terms of gene content and structure (Kortschak et al. 2003; Putnam et al. 2007) and have been shown to possess more homologs to human disease genes than traditional invertebrate models such as *Drosophila* and *C. elegans* (Maxwell et al. 2014). Thus, cnidarians are exceptional model systems for studying the mechanisms underlying pluripotency and lineage commitment, findings that can be applied within broader biological and biomedical contexts.

The colonial hydrozoan *Podocoryna* is an interesting cnidarian that has a complex life cycle that includes both a sessile polyp and free-swimming medusa (jellyfish) stage. The medusa stage plays a critical role in *Podocoryna*’s life cycle given its primary role in sexual reproduction, producing gametes that are released into the water column for fertilization. Once fertilized, a planula larva is formed that settles on a suitable substrate and develops into a new polyp, which then asexually produces a colony of polyps that bud new medusae. The medusae can also drift and disperse over a broader area, providing a mechanism for colonizing new habitats and increasing genetic diversity (Sanders and Cartwright 2015). *Podocoryna* has a proven history as a tractable laboratory model, due in large part to its ease of cultivation, its rapid life cycle, and its transparent nature, which facilitates live imaging and detailed observation (Braverman 1962; Braverman 1971). This experimental flexibility has been uniquely valuable for investigating questions related to coloniality, development, and regeneration (Schmid and Alder 1984; Weber et al. 1987; Aerne et al. 1995; Gröger and Schmid 2000; Blackstone and Buss 1992).

Evidence from experimental studies indicate that the development of the medusae involves a combination of conserved genetic pathways shared with bilaterians, as well as medusozoan-specific factors that are involved in the unique aspects of medusa formation, such as the development of the medusa bell, the arrangement of tentacles, and specialized muscle and nerve structures (Steinmetz et al. 2012; Leclere and Rottinger 2016; Travert et al. 2023).

Importantly, many of the cell types present in the medusa, including those comprising the striated muscle, exhibit a degree of ultrastructural complexity that is comparable to that found in bilaterian animals (Jahnel et al. 2014; Tanaka et al. 2018; Brunet et al. 2016). This mix of genomic novelty and conservation in an organism capable of whole-body regeneration positions *Podocoryna* as a compelling system for understanding the evolution of complex cell types.

*Podocoryna* is also proving to be a valuable model for the study of allorecognition – the ability to recognize self from non-self. Similar to its relative *Hydractinia*, it possesses a highly polymorphic allorecognition system that it uses to both recognize conspecifics and successfully compete for space in the marine environment (Tardent and Jauch 1983). Consistent with previous observations in *Hydractinia* (Huene et al. 2022), the *Podocoryna* allorecognition complex (ARC) is comprised of numerous highly duplicated allorecognition genes, with preliminary studies indicating significant homology between these genes across cnidarians as well as overall syntenic conservation throughout the phylum (manuscript in preparation). These observations suggest an evolutionary throughline between the ARC and similar regions responsible for allorecognition in other organisms, as well as with the human major histocompatibility complex (Ohta et al. 2019; Flajnik and Kasahara 2010).

Here, we present the first chromosome-level assembly of *Podocoryna americana* (Mayer, 1910), representing one of the most highly complete cnidarian genome assemblies to date. Our assembly comprises 327.2 Mb across 17 chromosome-scale scaffolds and 15 unplaced contigs. We use PacBio Iso-Seq, Illumina RNA-seq, and protein data to predict the presence of 19,085 protein-coding genes in the assembly and document numerous gene families with important roles in stem cell maintenance, striated muscle development and function, innate immunity, and allorecognition. Ortholog identification and macrosynteny analysis suggests that two chromosomal fission events have occurred in *P. americana* since its divergence from *Hydractinia*. Together, our chromosome-level genome assembly of *P. americana* will be an invaluable resource for the advancing the study of metazoan stem cell biology, pluripotency, development, and regeneration.

## Materials and methods

### Biological sample collection and animal husbandry

*Podocoryna americana* (Mayer, 1910) (NCBI: txid589949) is found on gastropod shells inhabited by hermit crabs along the east coast of North America. The parents of the genome strain were originally collected by the laboratory of Leo Buss (Yale University). The female parent strain (PA001) was collected at Old Quarry Harbor (Guilford, CT) and the male parent strain (PA002) was collected at Lighthouse Point (New Haven, CT) (Table 1). To produce the genome strain individual, 20 adult female (PA001) and male (PA002) medusae were placed together in a glass bowl for spawning and fertilization. Larvae were allowed to settle on microscope slides and primary polyps were fed *Artemia* nauplii starting at 72 hours after settlement. Two weeks after larval settlement, the slowest-growing colonies were removed to retain only one colony per slide. Two months after larval settlement, the mature colonies were sexed, and a male colony was selected to generate the reference genome (PA003). See the supplementary materials sections *Origin of genome parent strains* and *Production of the genome strain* for details.

**Table 1:** ***Podocoryna americana* sample origin details.**

| BioSample Accession | Name | Collection Site | Latitude, Longitude DMS | Sex |
| --- | --- | --- | --- | --- |
| SAMN51402013 | PA001 | Old Quarry Harbor, Guilford, CT | 41° 20' 16" N, 72° 10' 59" W | F |
| SAMN51402014 | PA002 | Lighthouse Point, New Haven, CT | 41° 14' 50" N, 72° 54' 13" W | M |
| SAMN51402012 | PA003 | F1 from University of Kansas | N/A | M |

### Sequencing library preparation and sequencing

For high molecular weight genomic DNA extraction, 200 non-reproductive polyps from the male genome strain individual (PA003) were dissected after one week of incubation in penicillin (50 u/mL)-streptomycin (50 µg/mL) (ThermoFisher Scientific, #15140122) and four days of starvation. For high molecular weight RNA extraction, 200 medusae from the male genome strain (PA003) and 200 non-reproductive polyps from the female parent strain (PA001) were collected after four days of starvation.

In total, we constructed 12 Illumina and PacBio DNA and RNA sequencing libraries. Seven of these libraries were from the male genome strain individual (PA003), four libraries were from the female parent strain (PA001), and one was from the male parent strain (PA002). Briefly, we obtained 263x coverage of PacBio WGS CLR reads from genomic DNA libraries of PA003 that were sequenced on a single PacBio Sequel II SMRTCell and 226x coverage from three Illumina Hi-C libraries sequenced on the Illumina HiSeq 4000. Hi-C data was generated using a Phase Genomics (Seattle, WA) Proximo Hi-C 3.0 kit, which is a commercially available version of the

Hi-C protocol (Lieberman-Aiden et al. 2009). We also obtained 472x and 423x coverage of Illumina WGS reads from libraries of the female parent strain (PA001) and male parent strain (PA002), respectively. Finally, we obtained a total of 1.48 million Iso-Seq transcripts from three libraries from medusae of the male genome strain (PA003) and three libraries from polyps of the female parent strain (PA001) that were sequenced on the PacBio Sequel II platform.

Sequencing was performed at the NIH Intramural Sequencing Center (Rockville, MD) and by Phase Genomics (Seattle, WA). See the supplementary materials sections *HMW DNA extraction for PacBio and Illumina WGS and Hi-C* and *HMW RNA extraction for Iso-Seq* for details.

### Mitochondrial genome assembly and phylogeny

The complete mitochondrial genome of *P. americana* was obtained by aligning mitochondrial genes present on a partial mitochondrial sequence present in GenBank (LN901210.1) to the PacBio WGS CLR reads using command-line BLAST+ v2.15.0 using default parameters (Camacho et al. 2009). Reads with mitochondrial hits were retrieved and used to generate a *de novo* assembly with CANU (Koren et al. 2017) using default parameters and an estimated genome size of 15 kb. The resulting contig had a length of 15,439 bp and manual annotation of the putative mitochondrial genome was carried out as described for the *Hydractinia* genome project (Schnitzler et al. 2024).

To determine the phylogenetic position and taxonomic identity of the genome strain individual (PA003), we constructed a mitochondrial 16S rRNA gene tree using sequences from species in the genera *Podocoryna*, *Hydractinia*, *Clava*, and *Stylactaria* in the family Hydractiniidae (Miglietta et al. 2009; Kayal et al. 2015) and outgroup sequences from the genus *Stylaster* (Lindner et al. 2008) using IQ-TREE v2.3.515. For details, see the *Mitochondrial genome assembly and annotation* and *Mitochondrial marker gene phylogeny construction* sections of the Supplementary Materials.

### Genome size estimation

Illumina WGS reads from the female parent strain (PA001) and male parent strain (PA002) were trimmed using Cutadapt v4.0 (Martin 2011), and k-mers were counted from trimmed reads using jellyfish v2.3.1 (Marcais and Kingsford 2011) using the settings -C -s 10000000000 -k 21. Genome sizes for each parent strain were estimated independently using the k-mer count histograms on the GenomeScope2.0 web server (Ranallo-Benavidez et al. 2020).

### Genome assembly and scaffolding

The PacBio CLR data was initially assembled into a draft genome using the FALCON pb-assembly pipeline (Chin et al. 2016). Haplotigs were removed using Purge Haplotigs v1.1.2 (Roach et al. 2018). Hi-C reads were aligned to the contig assembly and the Phase Genomics’ Proximo Hi-C genome scaffolding platform was used to create chromosome-scale scaffolds (Bickhart et al. 2017). Approximately 20,000 separate Proximo runs were performed to optimize the number of scaffolds and scaffold construction to make the scaffolds as concordant with the observed Hi-C data as possible. Juicebox v0.7.17 (Rao et al. 2014; Durand et al. 2016) was used to manually correct scaffolding errors. For details, see the *Genome assembly* section of the Supplementary Materials.

### Contaminant filtering and scaffold selection

All 538 scaffolds in the preliminary genome assembly were screened for contamination using BlobToolKit (Challis et al. 2020), which filtered out contaminant scaffolds based on GC content, coverage of RNA-seq data from NCBI BioProject PRJNA245897 (Sanders and Cartwright 2015) and PRJNA744579 (Travert et al. 2023), and the taxonomy of best BLASTN hits to the NCBI nt database. Additional identification and removal of contaminant sequences in assembled contigs was performed using FCS-GX (Astashyn et al. 2024). This process identified 34 chromosome-scale scaffolds and 269 unplaced contigs with best hits in the phylum Cnidaria, with the 34 chromosome-scale scaffolds representing 17 pairs of collinear chromosomes. To generate a haploid assembly with the highest level of completeness, BUSCO v6.0.0 was used in genome mode to search each chromosome-scale scaffold for metazoan single-copy orthologs, and the scaffold with the greater number of complete single-copy orthologs from each pair was selected for inclusion in the haploid assembly. These 17 chromosome-scale scaffolds were then ordered and numbered according to size. Next, BUSCO was used to identify single-copy orthologs on the 269 unplaced cnidarian contigs, and the 15 contigs containing 22 single-copy orthologs that were uniquely absent in the chromosome-scale scaffolds were retained in the final assembly.

### Repeat annotation and masking

To identify and mask repetitive regions in the genome prior to gene model prediction, RepeatModeler v2.0.1 (Flynn et al. 2020) was used with default parameters to create a custom repeat library that was taken as input by RepeatMasker v4.1.7 (Tarailo-Graovac and Chen 2009) to scan the genome and soft-mask repetitive sequences. The output of RepeatMasker was summarized using the helper scripts from RepeatMasker v4.1.7.

### PacBio Iso-Seq transcriptome generation pipeline

We generated full-length transcripts using the PacBio Iso-Seq v3 pipeline (https://github.com/PacificBiosciences/IsoSeq) using PacBio SMRT sequencing data from male medusa and female polyp samples. Raw subreads were processed into circular consensus sequences (CCS) using ccs (https://github.com/PacificBiosciences/ccs) with a minimum read quality of 0.9 (--min-rq 0.9), tagged for primers, barcodes, polyA positions, and read completeness (isoseq tag), then refined to remove polyA tails and artificial concatemers (isoseq refine). The resulting full-length non-chimeric (FLNC) reads were clustered into high-quality consensus isoforms (isoseq cluster) that were aligned to the genome with pbmm2 (https://github.com/PacificBiosciences/pbmm2) and merged into a final set of non-redundant full-length transcript isoforms (isoseq collapse) for each sample type.

### Gene model prediction and functional annotation

For gene model prediction, the soft-masked assembly was used as input to the BRAKER3 pipeline (Gabriel et al. 2024) using PacBio Iso-Seq data, Illumina RNA-seq data from NCBI BioProject PRJNA245897 (Sanders and Cartwright 2015) and PRJNA744579 (Travert et al. 2023), and a protein database containing sequences of related cnidarians and OrthoDB v12 (Tegenfeldt et al. 2025) as supporting evidence (Supplementary Table 5).

Protein sequences from the predicted gene models were assessed for homology to sequences in the NCBI nonredundant (nr) database and Swiss-Prot database by performing DIAMOND BLASTP searches with an E-value cutoff of 10^−5^. Protein domain prediction was performed using InterProScan v5.72-103.0 (Jones et al. 2014) and functional annotation was performed using eggNOG-mapper v2 (Cantalapiedra et al. 2021), PANNZER2 (Toronen et al. 2018), and the KEGG automated annotation server (Moriya et al. 2007). Using ampir (Fingerhut et al. 2021), we identified putative antimicrobial peptides (AMPs) as sequences with >85% predicted probability of antimicrobial activity.

### Ortholog analysis

OrthoFinder v2.5.4 (Emms and Kelly 2019) was used to identify orthologous gene families (also called orthogroups) and generate comparative genomics statistics. Proteomes of cnidarians with high-quality reference genomes, including representatives from the classes Anthozoa (*Nematostella vectensis*, *Exaiptasia diaphana*, *Acropora millepora*, *Orbicella faveolata*, *Pocillopora* spp., and *Xenia* spp.), Hydrozoa (*Hydractinia echinata*, *Hydractinia symbiolongicarpus*, *Clytia hemisphaerica*, *Hydra vulgaris*, *Hydra viridissima*, and *Turritopsis dohrnii*), Scyphozoa (*Aurelia* spp., *Chrysaora quinquecirrha*, *Nemopilema nomurai*, *Rhopilema esculentum*, and *Sanderia malayensis*) and Cubozoa (*Morbakka virulenta*) were downloaded from NCBI RefSeq or other sources where available (Supplementary Table 6). Outgroup proteomes included representatives from Bilateria (*Branchiostoma lanceolatum*, *Asterias rubens*, *Saccoglossus kowalevskii*, *Capitella teleta*, *Phoronis australis*, and *Pecten maximus*), Placozoa (*Trichoplax adhaerens*), Porifera (*Amphimedon queenslandica*, *Ephydatia muelleri*, and *Halichondria panicea*) and Ctenophora (*Hormiphora californensis*, *Bolinopsis microptera*, and *Mnemiopsis leidyii*) (Supplementary Table 6). A total of 33 proteomes were used in the analysis and were filtered to retain only the longest isoform for each gene model, and any mitochondrial sequences were removed before input into OrthoFinder v2.5.4 with default settings. A presence/absence table of orthogroups was created to visualize the set intersections of orthogroups shared among different taxa using UpSetR (Conway et al. 2017).

To estimate the evolutionary dynamics of orthogroup expansions and contractions in *Podocoryna*, a subset of 26 cnidarian and bilaterian taxa from the OrthoFinder analysis were used for analysis with CAFE 5 (Mendes et al. 2021), with the sponge *Halichondria panicea* as the outgroup. A species tree for the 26 selected taxa was constructed using 73 single-copy orthologs identified by OrthoFinder. Single-copy ortholog protein sequences were individually aligned using MAFFT v7.52614 (Katoh and Standley 2013) in L-INS-i mode and then trimmed using ClipKIT (Steenwyk et al. 2020) in the default “smart-gap” mode. The concatenated alignment was used for maximum likelihood phylogenetic analysis in IQ-TREE v2.3.515 with automatic model selection (-m MFP) (Kalyaanamoorthy et al. 2017), ultrafast bootstrapping (-B 1000), and SH-like approximate likelihood ratio test (-alrt 1000) enabled (Hoang et al. 2018; Anisimova et al. 2011). Divergence date estimates for major nodes on the tree were calculated using r8s v1.81 (Sanderson 2003) with node constraints specified for the divergence of Anthozoa and Medusozoa (570 Mya) and Hydrozoa (500 Mya). The time-calibrated tree was used together with the table of orthogroup sizes for each taxon as input to CAFE 5 to estimate gene copy numbers in internal nodes of the tree and identify branches with significant orthogroup expansions or contractions. As large orthogroups can impede parameter estimation in CAFE, orthogroups with counts >100 in any taxon were excluded. After testing several parameter sets, a Poisson distribution was used for the root frequency distribution (-p) and two gamma rate categories were used (-k 2). Orthogroups that were identified as significantly expanded or contracted in *P. americana* were extracted and tested for gene ontology (GO) term enrichment with topGO v2.58.0 (Alexa et al. 2006) using a database of GO terms assigned to the *P. americana* genome by eggNOG-mapper.

### Synteny analysis

The macrosyntR package (El Hilali and Copley 2023) was used to analyze macrosynteny between *Podocoryna* and cnidarian relatives for which chromosome-scale genomes have been generated, including the hydrozoans *Hydractinia symbiolongicarpus* and *Hydra vulgaris*, the scyphozoan *Rhopilema esculentum*, and the anthozoan *Nematostella vectensis*. For each species’ genome, gene model annotation was obtained from NCBI RefSeq and used to generate the required input BED files. macrosyntR was then used to perform pairwise synteny comparisons between *Podocoryna* and each of the other species based on the lists of single copy orthologs identified between each species. Finally, comparisons across all cnidarian genomes were completed to identify conserved gene linkage groups and chromosome-scale macrosynteny to infer the ancestral chromosome configuration and structural changes that have occurred during cnidarian evolution.

*Identification of candidate* Podocoryna *allorecognition genes (ALRs)*

Putative *P. americana* allorecognition genes (ALRs) were identified based on evidence of sequence homology and protein structural similarity. The predicted *P. americana* proteome was compared to a database of known *H. symbiolongicarpus* ALRs (Huene et al. 2022) using BLASTP and proteins with reciprocal minimum bitscore >50 were retained. Next, putative ALRs were required to have a predicted signal peptide (Teufel et al. 2022; Nielsen et al. 2024) and matches to published *H. symbiolongicarpus* ALRs with a TM-score >0.5 according to foldseek (van Kempen et al. 2024) and a root mean square deviation <2 angstroms according to DALI (Holm 2020).

## Results and discussion

### Chromosome-level assembly of P. americana (PodAmV1.0)

High molecular weight genomic DNA was extracted from a male *P. americana* colony (PA003, Table 1), and a total of 9,252,452 PacBio WGS CLR reads (263x coverage) were generated with a N50 read length of 13,187 bp (Supplementary Table 1), as well as a total of 247,898,772 Illumina Hi-C sequencing read pairs (Supplementary Table 1). Illumina WGS of the female parent strain (PA001) and male parent strain (PA002) yielded a total of 583,028,031 sequencing read pairs. High molecular weight RNA extracted from the genome strain (PA003) and its female parent strain (PA001) yielded a total of 861,700,001 PacBio Iso-Seq reads (Supplementary Table 1) and resulted in 57,422 non-redundant full-length transcripts after processing with the Iso-Seq v3 pipeline (Supplementary Table 2). The raw data are available on NCBI under BioProject PRJNA1329265 and details for each data type are available in Supplementary Table 1.

To determine the ploidy and estimate the genome size of *P. americana*, we computed k-mer spectra from the female and male parent strain Illumina WGS libraries. Both libraries had two major k-mer peaks, with a lower-coverage peak that was larger than a higher-coverage peak (Supplementary Figure 1). This pattern is consistent with the k-mer spectra of other highly heterozygous diploid organisms (Ranallo-Benavidez et al. 2020). Based on the k-mer spectra from the female and male parent strains, the predicted haploid genome size of *P. americana* was 325.2 Mb and 344.3 Mb, respectively (Supplementary Figure 1).

Next, we generated a *de novo* assembly of the *P. americana* genome from the PacBio WGS CLR reads using the FALCON pb-assembly pipeline. The preliminary assembly contained 1,350 contigs across 538 scaffolds with a total length of 717.4 Mbp. The scaffold N50 was 19.0 Mbp, the scaffold L50 was 17, and the total GC content was 37.26%. Contaminant filtering using BlobToolKit identified four scaffolds and 235 unscaffolded contigs (totaling 69.4 Mbp) from the preliminary assembly as contaminants based on best BLASTN hits to the NCBI nt database, which included representatives of the phyla Bacteroidota, Actinomycetota, Planctomycetota, and Pseudomonadota (Supplementary Figure 2). Additional filtering with FCS-GX removed two contaminant sequences (totaling 140.5 kbp) from the ends of two scaffolds.

The scaffold selection and contaminant filtering process resulted in a final 327.1 Mb haploid assembly (PodAmV1.0) with 17 chromosome-scale scaffolds and 15 unplaced contigs with a scaffold N50 of 20.7 Mbp, scaffold L50 of 7, and total GC content of 35.57% (Figure 1B, Table 2, Supplementary Figures 3 – 5). The 17 longest chromosome-scale scaffolds represent 98.0% of the assembly, ranging in size from 27.4 to 8.9 Mbp. BUSCO searches in genome mode detected 864 complete metazoan BUSCOs out of a total of 954 (90.6%), with 823 existing as single-copy (86.3%) and 41 as duplicates (4.3%) (Supplementary Table 3).

**Figure 1:**
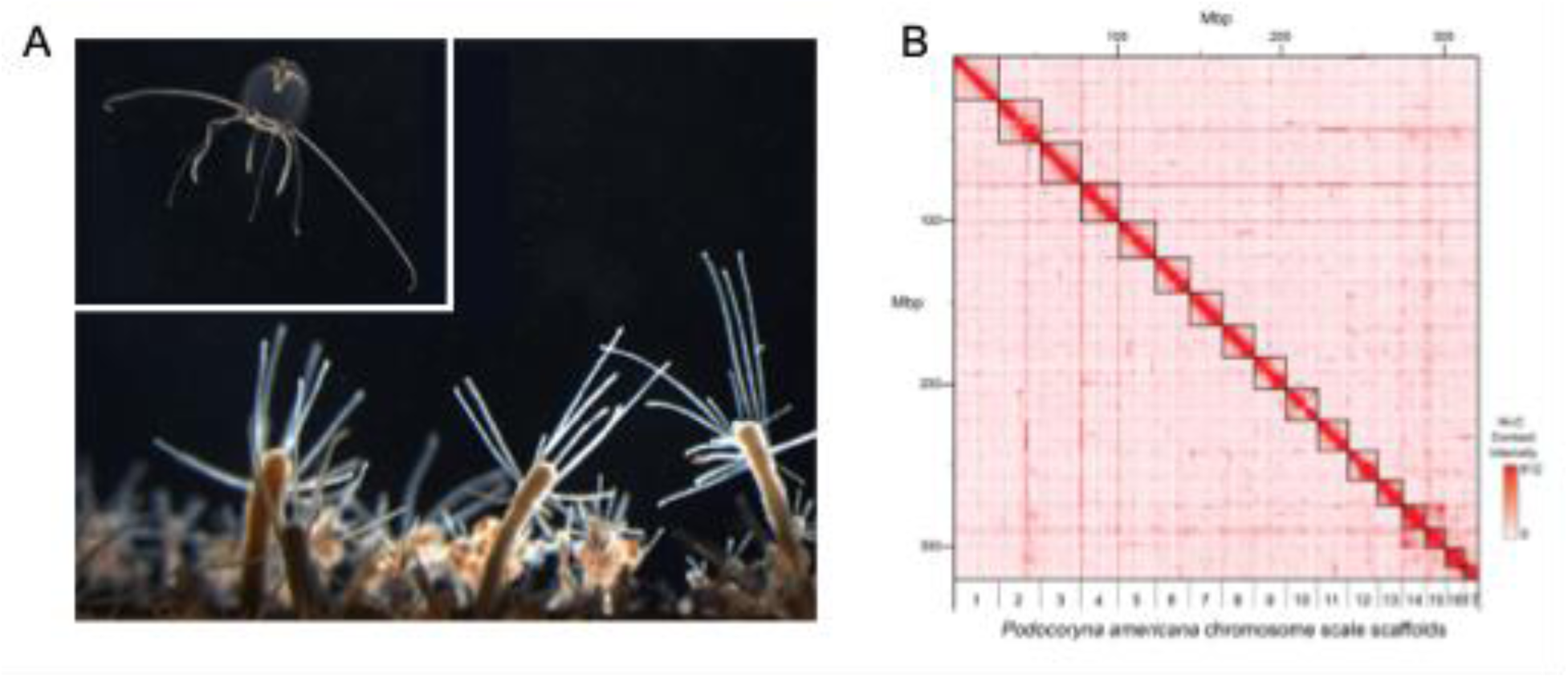
*P. americana* chromosome-level genome assembly. (A) Photographs of *P. americana* adult medusa stage (top) and colony stage consisting of feeding polyps and reproductive budding polyps (bottom). (B) Hi-C contact map for the chromosome-scale scaffolds in the *P. americana* genome assembly (PodAmV1.0).

**Table 2:**
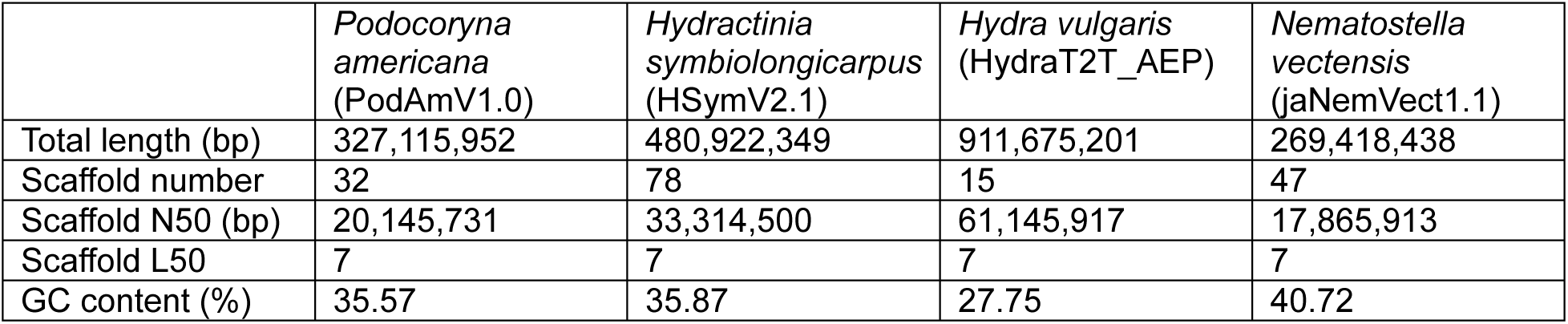
Comparison of Podocoryna americana genome assembly statistics with model cnidarians Hydractinia symbiolongicarpus, Hydra vulgaris, and Nematostella vectensis.

### Mitochondrial genome and phylogeny

We assembled the mitochondrial genome of the *P. americana* genome strain (PA003) and observed the typical mtDNA organization as described in other hydrozoans. The *P. americana* mitochondrial genome was identified as a linear 15,439 bp contig containing a set of 17 genes that includes the small and large ribosomal genes, one methionine and one tryptophan transfer RNA gene, and 13 energy pathway proteins (Supplementary Figure 6).

Similar to *H. symbiolongicarpus* and *H. echinata*, the mitochondrial genome of *P. americana* is comprised of a single linear chromosome that starts with an inverted repeat (IR), followed by a small pseudogene version of Cox1 and the large ribosomal (16S/RNL) subunit, both in a reverse transcriptional orientation. Replication and transcriptional orientation switches at the point of origin (Ori), identifiable by abrupt changes in DNA composition and stem-loop configurations containing T-rich loops. After Ori, the remainder of the *Podocoryna* mitochondrial genome is comprised of the following genes in order, as illustrated in Supplementary Figure 6: Cox2, tRNA-TRP, Atp8, Atp6, Cox3, tRNA-Met, Nad2, Nad5, RNS, Nad6, Nad3, Nad4L, Nad1, Nad4, Cob, Cox1, and finally, the capping IR. The IRs form G-rich loops that likely protect the ends of these linear mitochondrial chromosomes in the absence of telomeric sequences.

The mitochondrial 16S rRNA gene tree resulted in a trimmed alignment that was 605 bp in length, and ML phylogenetic analysis recovered the *Podocoryna* genome strain in a clade containing specimens identified as *P. americana* (Mayer, 1910) that were also originally collected from the North Atlantic coasts of the United States (Supplementary Figure 7).

### Distribution of repetitive elements and other genomic features

The *P. americana* genome assembly (PodAmV1.0) contains 156 Mbp (47.68%) of repetitive sequences, with each chromosome-scale scaffold having between 12.5 and 4.7 Mbp of repetitive sequences. The repetitive sequences are comprised of total interspersed repeats (45.14%), simple repeats (0.65%), and others (Supplementary Table 4). The total interspersed repeats are further divided into retroelements (5.8%), DNA transposons (3.64%), rolling circles (1.05%), and unknown repetitive sequences (35.7%), which are the most prevalent of all the repetitive sequences (Supplementary Table 4). Among the retroelements, LINE sequences (4.63%) were the most abundant, and among the DNA transposons, Maverick (0.75%) and hobo-Activator (0.41%) were the top two elements (Supplementary Table 4). Analysis of the *P. americana* repeat landscape suggests a continuous history of repeat expansion with a relatively recent peak of maximum repeat expansion activity (Figure 2).

**Figure 2:**
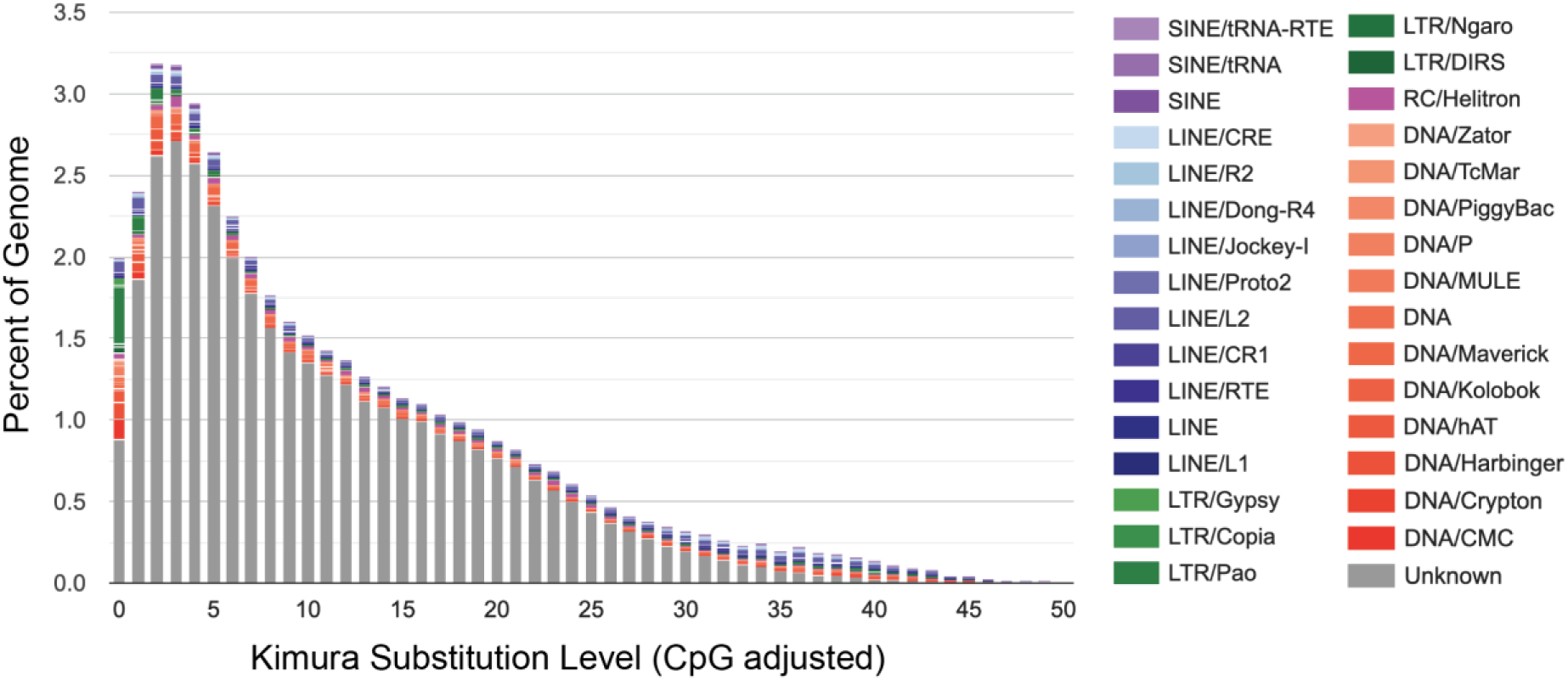
Interspersed repeat landscape of *P. americana*. The x-axis represents the level of Kimura substitution for repeat elements from the consensus sequences (relative age), while the y-axis represents the relative abundance of each repeat family in the genome. Ancient active repeats are placed on the right side of the graph and recently active repeats are on the left.

### Gene prediction and functional annotation

The BRAKER3 gene prediction pipeline yielded 19,085 protein-coding gene models with 22,865 predicted transcripts. The predicted protein-coding genes had a mean gene body length of 6,494 bp, a mean coding sequence (CDS) length of 1,872 bp, and a mean intergenic region length of 1,609 bp. On average, there were 1.20 transcripts, 8.78 exons, and 7.58 introns per protein-coding gene. In the 22,685 predicted proteins, we detected 870 complete metazoan BUSCOs out of a total of 954 (91.2%), with 730 existing as single-copy (76.5%) and 140 as duplicates (14.7%).

Of the 19,085 predicted protein-coding genes in our assembly, 17,269 protein-coding genes (90.5%) had significant homology (E-value <10^−5^) to sequences in the NCBI nonredundant database (nr), and 9,819 protein-coding genes (51.4%) had significant homology to sequences in the SwissProt database. A total of 18,077 protein-coding genes (94.7%) were functionally annotated using InterProScan, with 14,918 protein-coding genes (78.1%) having InterPro domain accessions and 13,483 protein-coding genes (70.6%) having PFAM protein domains. In addition, 10,078 protein-coding genes (52.8%) were functionally annotated using eggNOG-mapper and 7,095 protein-coding genes (37.2%) were functionally annotated using PANNZER2. Lastly, a genome-wide scan using ampir (Fingerhut et al. 2021) identified 162 protein-coding genes (0.8%) with putative antimicrobial activity. Overall, 18,679 protein-coding genes (97.9%) had functional annotation supported by one of more of the above databases (Figure 3).

**Figure 3:**
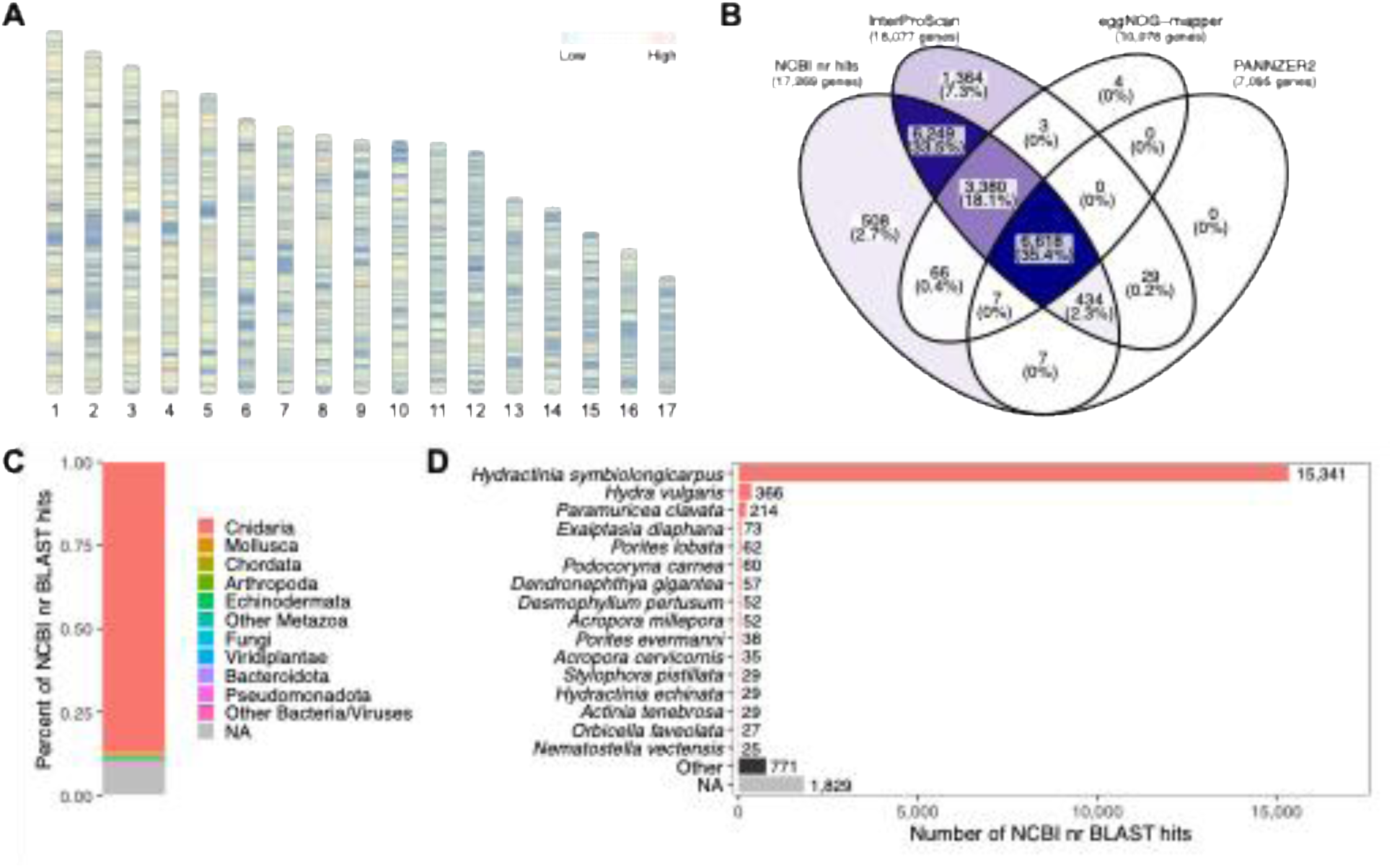
*P. americana* genome assembly functional annotation summary. (A) Ideogram displaying the density of predicted gene models across the 17 chromosome-scale scaffolds in 10 kbp windows, (B) Venn diagram depicting the number of predicted gene models receiving functional annotation from the NCBI nr, InterProScan, eggNOG-mapper, and PANNZER2 databases, (C) stacked bar plot of proportion of gene models with homologous protein sequences in the NCBI nr database by phylum, and (D) bar plot of top 16 species with homologous protein sequences in the NCBI nr database, other species and sequences without hits are shown at the bottom.

Notably, there were 60 protein-coding genes with best hits to *Podocoryna carnea* sequences present in NCBI GenBank that were previously characterized in early molecular cloning studies. These include the key homeodomain transcription factors *Otx* (Muller et al. 1999), *Gsx* (Yanze et al. 2001), and *Msx* (Galle et al. 2005), stem cell-related genes *piwi* (Seipel et al. 2004) and *nanos* (Torras et al. 2004), and genes coding for muscle structural proteins such as actin (Aerne et al. 1993), myosin heavy chain (Schuchert et al. 1993), and tropomyosin (Baader et al. 1993; Groger et al. 1999), among others.

### Ortholog identification, divergence time estimation, and gene family evolution analysis

A total of 741,387 protein sequences from 33 proteomes were used as input to OrthoFinder, which assigned 698,312 genes (94.2% of total) to 43,015 orthogroups. We identified 2,992 orthogroups (7.0%) shared among all cnidarians, with 3,432 orthogroups (8.0%) shared among medusozoans and 5,777 orthogroups (13.4%) shared among hydrozoans. A total of 8,224 orthogroups (19.1%) were shared across the family Hydractiniidae, and 353 orthogroups were identified as unique to the family Hydractiniidae. Of the 19,085 protein-coding genes in the *Podocoryna* genome, 652 were identified as *Podocoryna*-specific (3.4%) and were contained in 208 *Podocoryna*-specific orthogroups.

The concatenated alignment consisted of 73 single-copy orthologs shared across 26 species, comprising 45,491 amino acids with 34,702 variable sites, 10,121 parsimony-informative sites and 25.3% gaps (Figure 4). Divergence date estimation with r8s yielded an estimate of approximately 127 million years of divergence between *P. americana* and *H. symbiolongicarpus*, with the remaining estimates similar to those reported in Schnitzler et al. (2024). For example, the divergence time between *H. echinata* and *H. symbiolongicarpus* was estimated at 13.8 million years, while Schnitzler et al. (2024) estimated 19.1 million years of divergence between these sister species. The divergence time estimates for other hydrozoan, scyphozoan, and anthozoan clades were also similar between the datasets.

**Figure 4:**
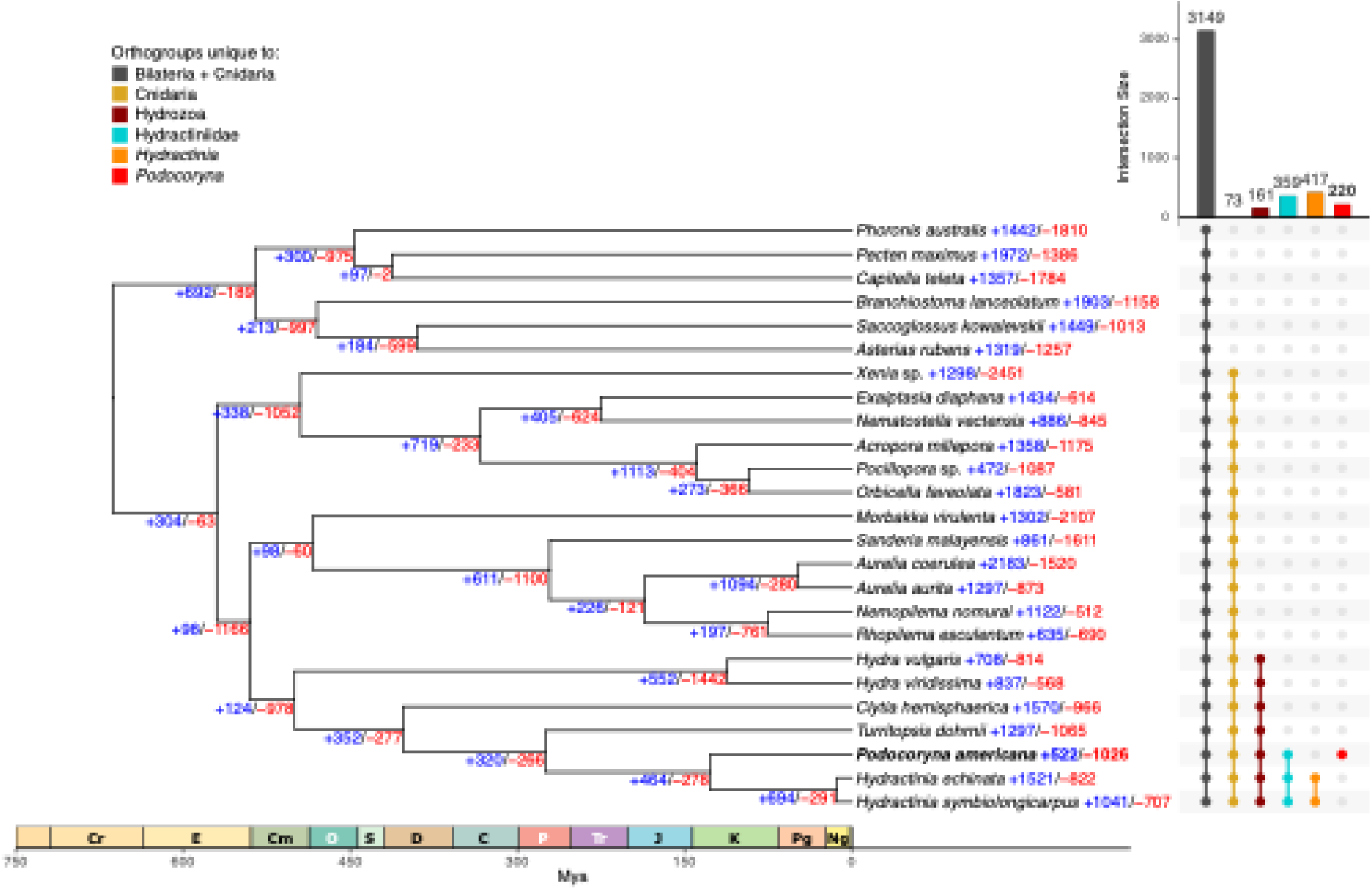
Conservation of *P. americana* gene content with cnidarian species revealed by OrthoFinder and CAFE analysis. UpSet plot displaying the number of shared orthogroups among selected taxonomic groups, including Bilateria and Cnidaria (dark grey), Phylum Cnidaria (gold), Class Hydrozoa (dark red), Family Hydractiniidae (turquoise), genus *Hydractinia* (orange), and *Podocoryna* (red). The colored circles below the vertical bar chart indicate those species that belong to each intersection group. The species tree constructed from an alignment of 73 single-copy orthologs (45,491 amino acid positions) was inferred by IQ-TREE2 and manually rooted with the sponge *Halichondria panicea* as an outgroup. The species tree was time-calibrated using r8s with priors for Hydrozoa (500 Mya) and Medusozoa (570 Mya). Node values depict the number of significant (p<0.05) gene family expansions (+, blue) and contractions (-, red) identified by CAFE 5.

A total of 36,915 orthogroups with counts <100 across all taxa were used for the CAFE model selection and parameter estimation steps. Gene family evolution analysis with CAFE identified 522 expanded and 1,026 contracted gene families in *P. americana* (Figure 4), with these gene families consisting of 3,010 and 1,408 genes, respectively. Expanded gene families in *P. americana* were significantly enriched for GO terms related to cell adhesion and morphogenesis, including regulation of extracellular matrix organization (GO:1903053, p_adj_ = 1.6×10^−6^) and cell-matrix adhesion (GO:0007160, p_adj_ = 1.8×10^−4^) as well as processes linked to vascular and immune-like responses such as acute inflammatory response (GO:0002526, p_adj_ = 4.4×10^−6^) and neutrophil chemotaxis (GO:0030593, p_adj_ = 5.3×10^−3^). Expanded gene families were also enriched for specific GO terms related to striated muscle structure and function including alpha-actinin binding (GO:0051393, p_adj_ = 1.1×10^−4^), spectrin-associated cytoskeleton (GO:0014731, p_adj_ = 1.7×10^−4^), positive regulation of sarcomere organization (GO:0060298, p_adj_ = 1.8×10^−4^), intercalated disc (GO:0014704, p_adj_ = 1.2×10^−3^), costamere (GO:0043034, p_adj_ = 4.6×10^−3^), and sarcolemma (GO:0042383, p_adj_ = 4.2×10^−2^), highlighting coordinated enrichment of key striated muscle structural complexes (Supplementary Table 7). Contracted gene families were enriched for genes involved in complex development and sensory/neural functions, including response to axon injury (GO:0048678, p_adj_ = 1.8×10^−6^), limb morphogenesis (GO:0035108, p_adj_ = 1.6×10^−4^), heart development (GO:0007507, p_adj_ = 8.3×10^−3^), and sensory perception of sound (GO:0007605, p_adj_ = 5.5×10^−4^), indicating preferential contraction of gene families associated with elaborate organ systems and regulatory pathways (Supplementary Table 7).

### Macrosynteny analysis reveals derived chromosomal fissions in Podocoryna

*Podocoryna* exhibits conserved chromosome macrosynteny with other chromosome-scale cnidarian genomes (Figure 5). Pairwise analysis of 8,114 single-copy orthologs shared between *Podocoryna* and its closest relative with a chromosome-level genome, *H. symbiolongicarpus*, revealed one-to-one macrosynteny between 13 pairs of chromosome-scale scaffolds despite extensive gene order rearrangement within each of the chromosomes. Notably, our analysis detected two chromosomal fission events where *Hydractinia* chromosomes 6 and 15 appear to have split, generating the pairs of *Podocoryna* chromosomes 14 and 15, and 16 and 17, respectively (Figure 5). These two chromosomal fissions appear responsible for the difference in chromosome number between *Hydractinia* (15) and *Podocoryna* (17); however, a chromosomal karyotype would be required to validate this result.

**Figure 5:**
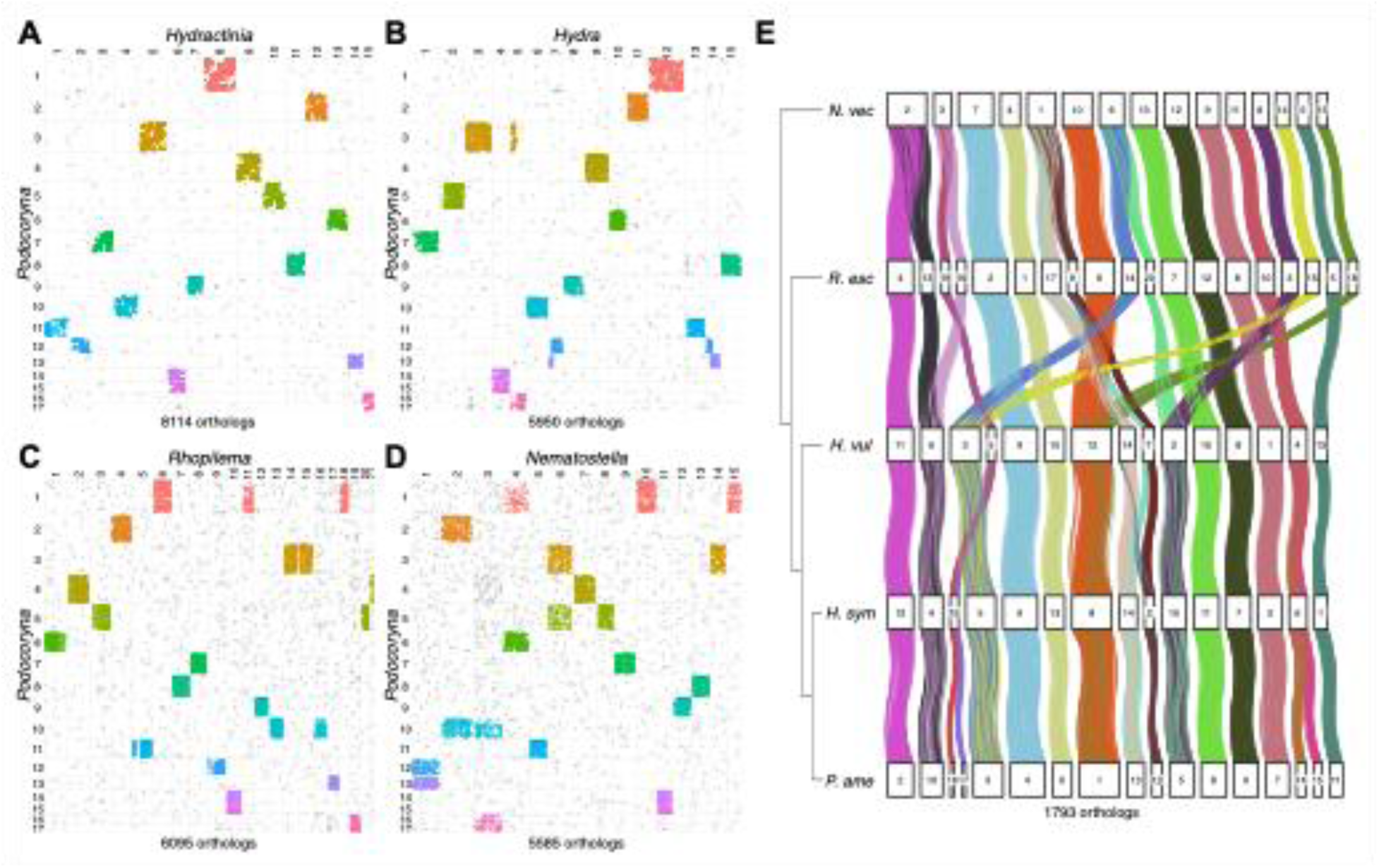
Chromosome-scale macrosynteny between *Podocoryna* and cnidarian model species. Oxford dot plots of single copy orthologs between *P. americana* and cnidarian model species (A) *Hydractinia symbiolongicarpus*, (B) *Hydra vulgaris* AEP strain, (C) *Rhopilema esculentum*, and (D) *Nematostella vectensis*. Chromosome-level scaffolds are ordered according to size along the x-axis and y-axis, and points representing single copy orthologs in linkage groups are colored according to *Podocoryna* chromosome number to highlight conserved chromosome scaffolds in pairwise species comparisons. Orthologs not found in linkage groups are colored grey. The number of single-copy orthologs identified on chromosome scaffolds in each pairwise species comparison is shown below the plot. (E) Ribbon diagram of chromosome-scale gene linkages across *Nematostella*, *Rhopilema*, *Hydra*, *Hydractinia*, and *Podocoryna* reveals the occurrence of chromosomal fission, translocation, fusion, and fusion-with-mixing events in the evolution of cnidarian chromosomes. Numbered boxes represent the chromosomes for each species, and ribbons connecting the boxes represent the coordinates of 1,793 single-copy orthologs contained within the conserved linkage groups across the five species. Ribbons are colored according to linkage groups.

In addition, the sizes of syntenic chromosomes varied between *Hydractinia* and *Podocoryna*, with *Podocoryna* chromosome 1 demonstrating synteny with *Hydractinia* chromosome 8, *Podocoryna* chromosome 2 with *Hydractinia* chromosome 12, and so on, demonstrating that changes in both chromosome size and gene order that have occurred in these hydrozoan lineages despite the continued maintenance of chromosome-scale gene linkages (Fig 5A-D).

Next, quintet analysis of *Podocoryna*, *Hydractinia*, *Hydra*, *Rhopilema*, and *Nematostella* based on 2,649 single-copy orthologs shared between all five species identified 1,793 orthologs that belonged to 23 conserved linkage groups (Figure 5E). Inspection of the macrosynteny ribbon diagram for these conserved linkage groups revealed that different chromosomal fusions, translocations, and gene order rearrangements that have occurred in hydrozoans and anthozoans relative to the ancestral cnidarian chromosomal configuration that is preserved in *Rhopilema* (Simakov et al. 2022). For example, despite *Nematostella*, *Hydra* and *Hydractinia* all having shared chromosome numbers of 15, at least four chromosomal fusion-with-mixing events have occurred in the hydrozoan common ancestor following the split with Anthozoa, whereas at least three different fusion-with-mixing events occurred in the anthozoan lineage (Figure 5E).

### Identification of candidate Hox cluster, sex determination, and allorecognition complex (ARC) regions

Our functional annotation identified 77 putative homeobox genes in the *Podocoryna* genome assembly, encoding 92 homeodomain-containing proteins. Of these, 36 genes were members of the Antennapedia (ANTP) class, including the two extended Hox genes *Eve* and *Mox*, 10 Hox/Hox-like genes, three ParaHox genes and 23 NK/NK-like genes (Supplementary Table 8). Several of the Hox and all three ParaHox genes were found to be distributed along two chromosomes, six of the NK genes along with the extended Hox gene *Mox* were distributed along a single chromosome and the NK gene *Msx* and *Eve* clustered on a third chromosome. Although macrosynteny and gene order and clusters were conserved between *Podocoryna* and *Hydractinia*, *Eve*, the Hox genes *Scox-l*, *Hox1*, *Pdx*, *Cnox2* and the NK gene *Tlx* were absent in the *Hydractinia* genome (Supplementary Figure 8). Interestingly, the gene *Tlx* has previously been shown to be lost convergently in multiple cnidarian lineages, including *Hydractinia* (Travert et al. 2023).

Macrosynteny analysis and comparison with the *Hydractinia symbiolongicarpus* genome suggests that the regions responsible for sex determination in *Podocoryna* should be located on chromosome 11, which is syntenic with *Hydractinia* chromosome 1 (Kon-Nanjo et al. 2023; Chen et al. 2023). Similarly, the *Podocoryna* allorecognition complex is located on chromosome 7, which is syntenic with *Hydractinia* chromosome 3 that contains the published allorecognition complex (Huene et al. 2022; Kon-Nanjo et al. 2023). The *Podocoryna* allorecognition complex on chromosome 7 contains 22 putative allorecognition genes (ALRs), however there are some putative ALRs that are located outside of this complex; for example, *Podocoryna* chromosome 4 contains 4 ALRs, and chromosomes 1, 6, 11, and 14 each have a single putative ALR.

## Conclusion

In summary, we have sequenced and generated a chromosome-level *de novo* assembly of the marine hydrozoan *Podocoryna americana* genome having a total size of 327.2 Mb and scaffold N50 of 20.1 Mb. We also recovered the *P. americana* mitochondrial genome as a 15,439 bp linear contig. Hi-C scaffolding resulted in 17 chromosome-scale scaffolds containing 98% of the nuclear genome assembly. We used Illumina RNA-seq, PacBio Iso-Seq, and protein data to predict 19,085 protein-coding genes in this assembly, and we generated functional annotation for 18,679 (97.9%) of these protein-coding genes. Our novel chromosome-level genome assembly for *P. americana* will be an invaluable resource for the study of cell type evolution, pluripotency, and regeneration in cnidarians and other diverse metazoan species.

## Data availability

Data generated for this study is available under NCBI BioProject PRJNA1329265. PacBio WGS CLR data and Illumina Hi-C data used for the genome assembly are available on the NCBI Sequencing Read Archive (SRA) under accessions SRR35425041 and SRR35425035, SRR35425036, and SRR35425040, respectively. PacBio Iso-Seq data are available via SRA accessions SRR354250329 - SRR35425034 and SRR35425037 - SRR35425039. Illumina RNA-seq data are available via NCBI BioProject PRJNA245897 and PRJNA744579. The *P. americana* genome assembly has been submitted to NCBI under submission ID SUB15946590 and will be released upon completion of processing and publication of this manuscript. All data analysis scripts are available at https://github.com/michaeltconnelly/podocoryna_genome.

## Author’s contributions

E.S.C., A.D.B. and P.C. conceived and designed the study, M.T. and P.C. performed animal husbandry and completed DNA and RNA extractions, E.S.C. and A.D.B. coordinated the genome and transcriptome sequencing, M.T.C. performed the genome assembly and annotation, and S.N.B., A.M.R, A.M.K., and R.Y. contributed to the bioinformatics analyses and interpretation of the data. E.S.C., M.T.C., and A.D.B. drafted the manuscript, and all authors have reviewed and approved the final version of the manuscript.

## Acknowledgments

We extend our gratitude to the NIH Intramural Sequencing Center (NISC) for their assistance with PacBio WGS CLR and Iso-Seq sequencing of the *P. americana* genome and transcriptome. We also thank the staff at Phase Genomics for their support generating Illumina Hi-C reads and the preliminary chromosome-scale genome scaffolds.

